# Computational modeling of human genetic variants in mice

**DOI:** 10.1101/2025.02.23.639784

**Authors:** Kexin Dong, Samuel I. Gould, Minghang Li, Francisco J. Sánchez Rivera

## Abstract

Mouse models represent a powerful platform to study genes and variants associated with human diseases. While genome editing technologies have increased the rate and precision of model development, predicting and installing specific types of mutations in mice that mimic the native human genetic context is complicated. Computational tools can identify and align orthologous wild-type genetic sequences from different species; however, predictive modeling and engineering of equivalent mouse variants that mirror the nucleotide and/or polypeptide change effects of human variants remains challenging. Here, we present H2M (human-to-mouse), a computational pipeline to analyze human genetic variation data to systematically model and predict the functional consequences of equivalent mouse variants. We show that H2M can integrate mouse-to-human and paralog-to-paralog variant mapping analyses with precision genome editing pipelines to devise strategies tailored to model specific variants in mice. We leveraged these analyses to establish a database containing > 3 million human-mouse equivalent mutation pairs, as well as *in silico*-designed base and prime editing libraries to engineer 4,944 recurrent variant pairs. Using H2M, we also found that predicted pathogenicity and immunogenicity scores were highly correlated between human-mouse variant pairs, suggesting that variants with similar sequence change effects may also exhibit broad interspecies functional conservation. Overall, H2M fills a gap in the field by establishing a robust and versatile computational framework to identify and model homologous variants across species while providing key experimental resources to augment functional genetics and precision medicine applications. The H2M database (including software package and documentation) can be accessed at https://human2mouse.com.

## Introduction

A central goal of human genetics is to learn how genetic variants impact the cellular and molecular phenotypes that underpin human diseases. Genetically-engineered mouse models (GEMMs) are powerful tools for these types of functional studies due to their extensive genetic homology with humans and their *in vivo* physiological relevance. In recent decades, the integration of molecular cloning and genetic engineering have helped further establish the utility of GEMMs to study disease-related variants and model genetic diseases like cancer^1–4^. Next-generation sequencing technologies, combined with CRISPR-based genome perturbation methods, are also being increasingly exploited in mice to generate large datasets and accelerate the understanding of human diseases^5–8^.

Species-specific genetic inconsistencies represent a major challenge that complicates the development and benchmarking of GEMMs for studying human genetic variation and accurately interpreting biological effects^9^. This challenge consists of at least three problems. First, the complex nonlinear mapping of gene orthologs makes it difficult to find orthologous murine loci that can also be engineered in the mouse genome. This includes scenarios like the absence of a murine orthologous gene or functional domain^10^ and a variable number of paralogs^11^. Second, the functional effect of altering the orthologous sequence may vary in different species depending on local sequence contexts. Here, we define NCE (nucleotide change effect) as the DNA-level modification induced by a mutation, and PCE (peptide change effect) as the protein-level change (i.e. amino acid change). These are important to distinguish because engineerable NCEs may produce distinct PCEs depending on how much variation there is in the local sequence context between the human and mouse genomes. This is particularly important for mutations at loci where mice and humans have different splicing donor sequences or variable codon usage for the same exon or amino acid^12,13^. Third, variants with the same NCE and/or PCE at conserved sites do not necessarily play the same functional role in humans and mice in part due to interspecies differences in protein-protein interactions^14^ and genetic regulatory networks^15^, among others. Thus, a deep understanding of human variants of interest is critical to develop biologically-meaningful GEMMs and accurately interpret relevant mouse genetic data^16^. This is especially relevant in the current genome editing era, where one can conceivably engineer and interrogate millions of genetic variants at will.

While existing genetic resources and computational tools can help, there is an unmet demand for integrative, high-throughput tools that act as comprehensive dictionaries of cross-species genetic variants to model and engineer mutations with identical sequence and/or functional changes. Humans and mice have well-annotated reference genomes that allow mapping and alignment of gene orthologs^10,17–20^. Large public collections of genetic variants are also available for both species^21–24^. In terms of functional analysis, multiple tools are designed to search genetic regulatory networks to predict the pathogenic effects of mutations in both species^15,25,26^. However, no existing tools can provide *de novo* predictions of equivalent human-mouse NCEs and PCEs, especially if they have not been observed or studied before in either species. Furthermore, the scientific democratization of emerging high-throughput precision genome engineering methods^27^ for functional studies of orthologous mouse variants requires automated, user-friendly prediction tools that avoid the need for error-prone manual verification across multiple online resources. The results should also be standardized to perform other downstream analyses, including guide RNA design and functional prediction of pathogenic effects.

With the above goals in mind, we developed H2M (human-to-mouse), a computational pipeline that processes human genetic variation data to model and predict the functional consequences of equivalent mouse variants, as well as devise strategies for precision engineering and analysis of corresponding mutations in mice. H2M is robust and versatile; for instance, it can take as input genetic variants in flexible formats compatible with well-established public databases to systematically identify, model, and visualize orthologous variants across thousands of mutations. Importantly, while we showcase its utility for human-to-mouse and mouse-to-human analyses in this study, H2M is compatible with any organism that has a sequenced reference genome. We envision that H2M will enable facile modeling and functional characterization of human genetic variants in mice and other model organisms, allowing the comprehensive dissection of the cellular and molecular functions of the growing compendium of human genetic diversity.

## Results

### H2M allows precise modeling of human genetic variants in the mouse genome

To model human genetic variants in the mouse genome, the H2M workflow involves four main steps: (1) querying the orthologous gene, (2) aligning wild-type transcripts or peptides, (3) simulating the mutation, and (4) checking and modeling its functional effects (**Fig. 1a, b, Extended Data** Fig. 1).

**Figure 1.**
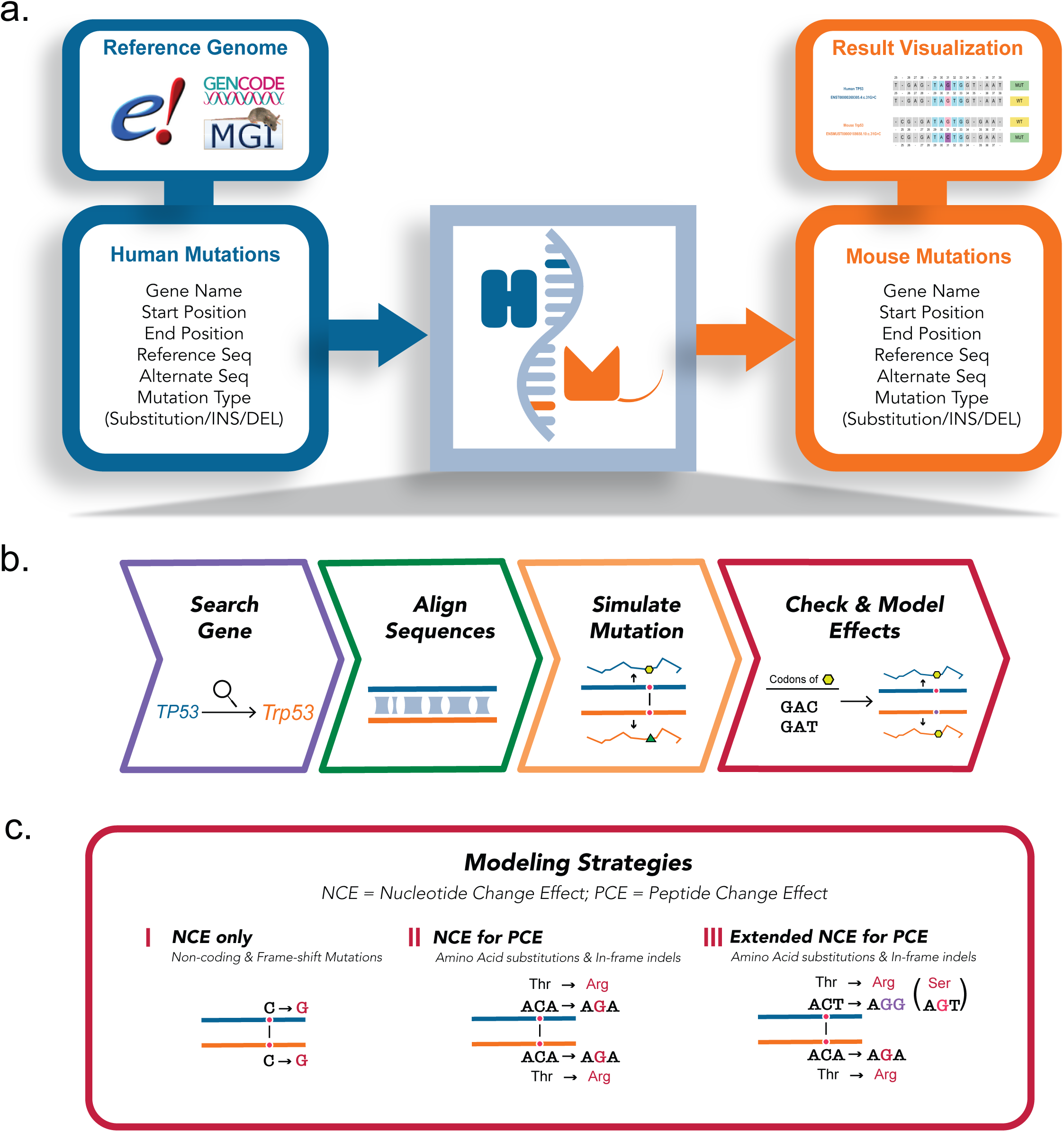
High-throughput computational modeling of human variants in the mouse genome. **a,** Schematic of the H2M pipeline. H2M takes as input human mutations, integrates data from databases including Ensembl, GENECODE, and MGI, and outputs and visualizes their murine counterparts in a high-throughput manner. INS = small insertion mutations, DEL = small deletion mutations. **b,** Four main steps of the H2M pipeline, (1) querying the orthologous gene, (2) aligning wild-type transcripts or peptides, (3) simulating mutation, and (4) checking and modeling its effect. **c,** H2M has different modeling effects depending on the specific sequence-change effect of the input. For non-coding and frame-shifting mutations, (I) NCE-only strategy is used to model the same DNA-level alteration. For amino acid substitutions, insertions and deletions, H2M takes either (II) NCE for PCE strategy, if the DNA mutation leads to the same amino acid change in both genomes, or (III) Extended NCE for PCE strategy, if a different DNA mutation is needed to model the target amino acid change. NCE = Nucleotide Change Effect, PCE = Peptide Change Effect.

Modeling equivalent genetic variants from different species requires the presence of homologous genes in each genome^16^. Approximately 1% of human genes do not have a mouse ortholog, and *vice versa*^10^. For example, the human gene *GPR32* has no mouse ortholog^28^. The Mouse Genome Database (MGD) and the Ensembl database allow for online queries of human-mouse orthology relationships^19,24,29,30^. Thus, we first pre-queried and integrated the lists of mouse and human homologs from these two sources and built them into the H2M package (**Supplementary Table 1**). H2M also provides an Application Programming Interface (API) based function for sending single-gene homology query requests to the Ensembl database, thereby ensuring data synchronization. Moreover, H2M is designed to aggregate both reference genomes and gene annotations from Ensembl and GENCODE databases^19,31^. The Ensembl Canonical Transcript ID — a single, representative transcript identified for each human or mouse gene — is also provided to inform the user’s choice of a proper transcript version that is used by default in batch process^19^ (**Supplementary Table 1**). After finding gene pairs, H2M retrieves complete sequences and all transcript versions for each gene. For a given transcript, H2M locates exons and introns, simulates RNA splicing, and obtains complete transcript sequences. H2M is compatible with any human and mouse reference genome chosen by the user, with optimal compatibility for GRCh37, GRCh38, and GRCm39. This also retains the potential for expansion to other species and experimental model organisms as they become available.

Once the genetic data is prepared using the above flexible interface, H2M proceeds to simulate, check, and model the functional effects of target gene mutations at both nucleotide and peptide levels. Mutations in poorly-conserved regions cannot be reliably modeled across humans and mice. To determine if mutations map to locally conserved regions, H2M aligns wild-type transcripts (for non-coding mutations) or peptide sequences (for coding-mutations) of human and mouse genes by using the Needleman-Wunsch algorithm. If the human mutation has a corresponding site in the mouse genome, H2M employs three main modeling strategies (**Fig. 1c**, **Table 1**). For all entries, H2M computes the same nucleotide change in the mouse transcript and outputs the NCE equivalent (Strategy #1: NCE-only Modeling, **Extended Data** Fig. 2a-c). Due to the fact that the same nucleotide alteration at corresponding human and mouse loci will not always result in the same amino acid, H2M also computes the effects of sequence changes at the protein level (PCE) for coding variants. To account for these potential differences, H2M also generates DNA-level changes that should produce the same protein-coding effect in both species. After simulating the same NCE in both genes and comparing the resulting amino acid alterations, H2M keeps the variant equivalent that mirrors both the NCE and PCE (Strategy #2: NCE for PCE Modeling, **Extended Data** Fig. 2d). Otherwise, H2M will try to provide extended PCE equivalents with different NCEs by leveraging codon redundancy (Strategy #3: Extended NCE for PCE Modeling, **Extended Data** Fig. 2e).

**Table 1.** H2M Modeling Strategies.

| Modeling Strategy | Mutation Type | Mutation Effects | Nucleotide Changes (Human) | Extended NCE for PCE Modeling Strategy |
| --- | --- | --- | --- | --- |
| (1) NCE-only Modeling | Non-coding | Intron;<br>Splice Region;<br>5'/3' Flank/UTR | SNP/DNP/ONP/<br>INS/DEL | N/A |
|  | Coding | Frame-shift/Nonstop | INS/DEL |  |
| (2) NCE for PCE Modeling<br>or<br>(3) Extended NCE for PCE Modeling | Coding | Missense/Silent | SNP/DNP/ONP | Same amino acid substitution(s) |
|  |  | In-Frame Insertion/Deletion | INS/DEL | Same amino acid insertions or deletions |
|  |  | Nonsense | SNP/DNP/ONP | Stop codons |
|  |  | Nonstop | SNP/DNP/ONP | All coding amino acid codons |

The output of H2M includes a wealth of standardized information that can be used for many different types of downstream analyses (**Table 2**). In addition to mutation coordinates and DNA-level sequence alterations in MAF format, H2M also provides transcript and protein-level sequence change effects using standard HGVS Nomenclature^32^.

**Table 2.**
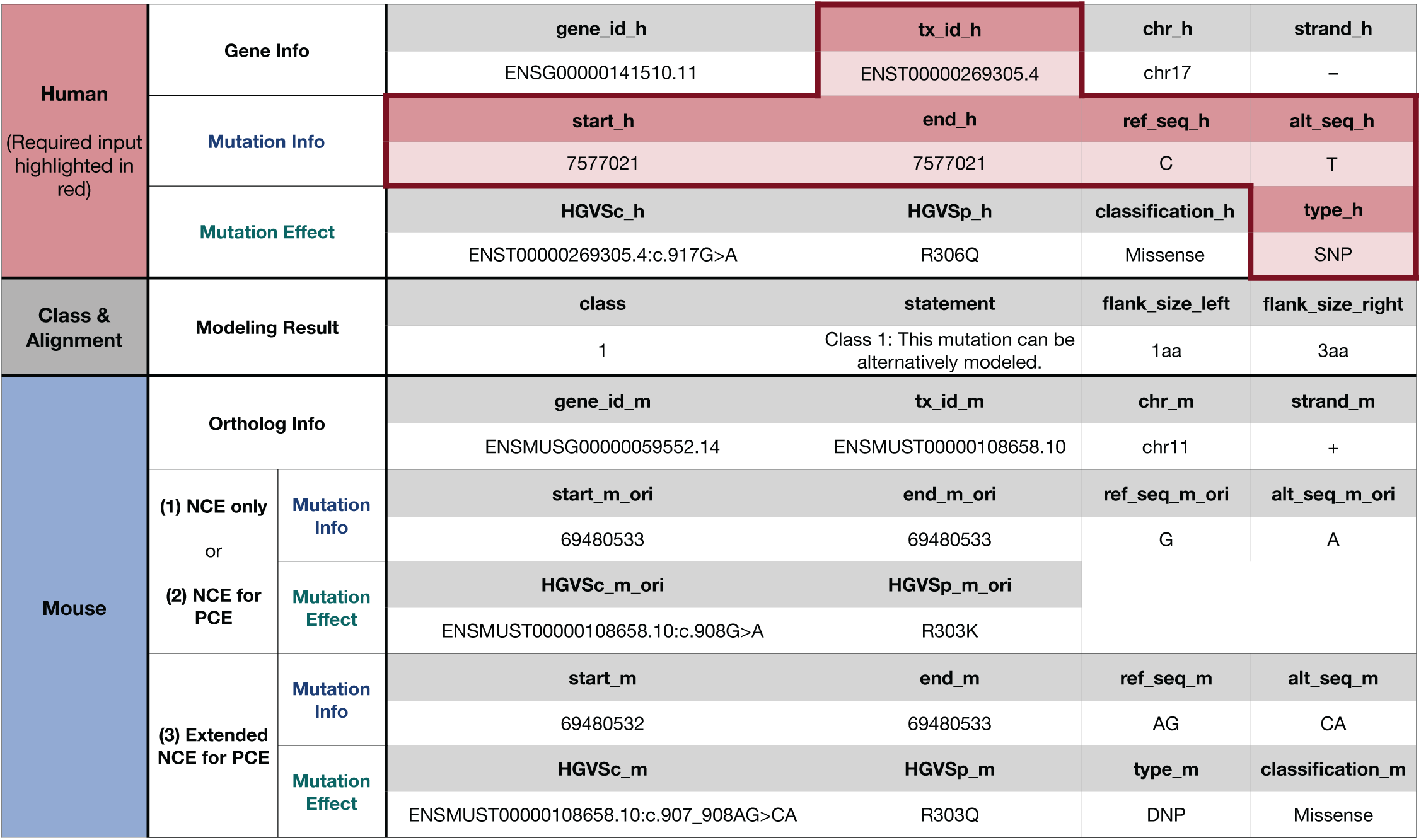
Representative H2M Output.

### H2M generates a mouse variant database from > 3 million human-to-mouse mutation mappings

Large-scale human genome sequencing studies have cataloged millions of germline and somatic mutations observed in humans. The development of a corresponding catalog of these mutations in mice would be of significant value for predicting the functional effects of these mutations and devising experimental strategies to establish new mouse models and interpret available experimental data from existing mouse models.

To this end, we first queried the AACR-GENIE, COSMIC, and ClinVar databases to retrieve human variants involving nucleotide substitutions and small insertions and deletions **(Fig. 2a)**. AACR-GENIE reports mutations based on clinical-sequencing data of cancer patients^23^, while COSMIC focuses on somatic mutations broadly observed in human cancer^33^. ClinVar is a public archive of human genetic variants, most of which are germline, along with information on their potential significance^34^. We then used H2M to filter this input to identify human-mouse gene-level orthologous relationships. As a result, 96% of the input human genes were mapped to their mouse orthologs, while 4% of the recorded genes were filtered out due to the lack of either a mouse ortholog or homologous relationship annotation (**Fig. 2b**). Most of the mappings were one-to-one, except for a handful of gene families with multiple paralogs, including the Pramel (preferentially expressed antigen in melanoma-like) and zinc finger protein (ZFP) families. We then used H2M to predict the murine equivalents and established the H2M Database (version 1), a dictionary encompassing 3,171,709 human-to-mouse mutation mappings (May 2024) (**Fig. 2a, Supplementary Table 2-3**).

**Figure 2.**
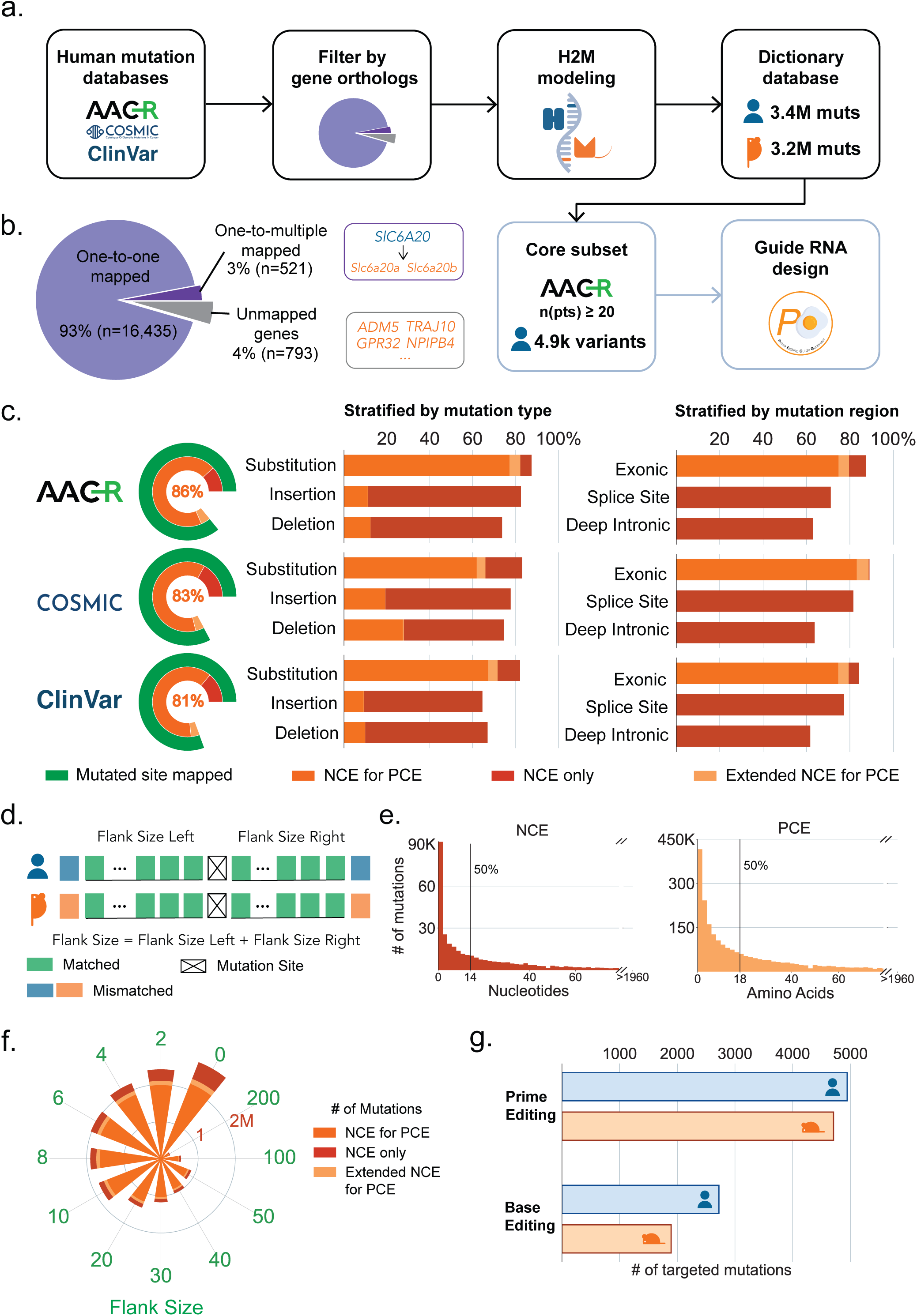
A human-to-mouse dictionary of clinically-observed genetic variants. **a,** Schematic of the H2M Database. We aim to generate a mutation database containing murine equivalents of human-observed genetic variants. Human variants are retrieved from AACR-GENIE, COSMIC and ClinVar Database and mapped to the mouse genome via H2M. Guide RNAs for precision genome editing, including prime editing and base editing, are designed for a selected subset of human mutations with high recurrent frequency as well as their murine equivalents. **b,** Pie chart visualizing the presence of mouse gene orthologs for human genes in the input human dataset. Most of the mutated human genes have a unique mouse homolog, and a few have multiple or none. **c,** The percentages of human mutations in the H2M Database that can be modeled in the mouse genome, stratified by the data source, modeling strategy, and regions and types of mutation. **d,** Schematic of the flank sequence for the mutation site. Flank size is defined as the combined length of consensus nucleotides (for non-coding variants) or peptides (for coding variants) on both sides of the mutated site. **e,** Distribution of flank sizes for all the human variants in H2M Database, split by NCE (left) for non-coding mutations and PCE (right) for coding mutations. **f,** The relationship between percentage of modelable mutations and flank size in the H2M Database. **g,** The number of mutations that are prime-editing and base-editing amenable in the selected subset of the H2M Database.

Remarkably, H2M predicts that >80% of human genetic variants can be modeled in the mouse genome (**Fig. 2c**). These variants are anatomically and functionally diverse, as H2M is able to model substitutions, insertions, and deletions. The majority of modelable variants fall under NCE-only and NCE-for-PCE categories, which can be modeled by introducing the same nucleotide-level mutations in an automated and robust manner. The NCE-only and NCE-for-PCE modeling categories, along with their corresponding mutations, represent one of the most highly robust applications of H2M to perform high-confidence cross-species modeling. As expected, we observed a slightly lower coverage for insertions and deletions compared to single or multi-nucleotide substitutions. When stratified by mutation regions, we found that a higher percentage of coding mutations can be modeled compared to those in non-coding regions, consistent with the higher sequence conservation in coding regions. Within non-coding regions, mutations in splice sites show a higher modeling prediction percentage than those present in deep intronic areas.

To take into account species-specific sequence differences surrounding a site corresponding to a variant of interest, we introduced a flexible parameter called “flank size”, defined as the combined length of consensus nucleotides (for non-coding variants) or amino acids (for coding variants) on either side of the mutated site (**Fig. 2d**). In the H2M database, 50% of coding mutations have a flank size of 18 or fewer amino acids, and 50% of non-coding mutations have a flank size of 14 or fewer nucleotides (**Fig. 2e**). When locating the mutation site, H2M can filter out mutations below a specific flank size provided by the user. As expected, the percentage of variants identified as modelable by H2M reduces as the size of the flank expands, as it restricts engineerable mutations to regions with higher sequence homology (**Fig. 2f**).

### H2M integration with precision genome editing to enable cross-species functional genetic analysis

Recent advances in genome engineering, including the development of precision genome editing technologies like base editing and prime editing, are enabling researchers to precisely and efficiently engineer and study mutations of interest within their native genetic environments^27^. Still, comparative functional genetic and genomic studies remain challenging in large part due to lack of computational tools that enable accurate and systematic identification and design of genome editing strategies that faithfully mirror cross-species genetic changes. In addition to providing a list of engineerable mutations across species, it is also essential to identify guide RNAs with predicted on-target and/or off-target characteristics that could be used for single or multiplexed variant studies. To test whether H2M could also do this, we selected a subset of 4,944 recurrent cancer-associated human mutations along with their mouse counterparts and utilized PEGG (Prime Editing Guide Generator) to design guide RNAs for base editing and prime editing (pegRNAs) (**Fig. 2a**)^35^. These analyses allowed us to generate a first-of-its-kind database containing 24,680 unique base editing gRNAs targeting 4,612 mutations (2,720 human mutations and 1,892 mouse mutations), and 48,255 unique pegRNAs targeting 9,651 mutations (4,944 human mutations and 4,707 mouse mutations) across the human and mouse genomes (**Fig. 2g, Supplementary Table 4-5**). Recent advances in precision genome editors, such as NCN-context-dependent CBEs^36^ and PAM-flexible variants like SpCas9-NG^37^, have greatly expanded targeting possibilities. To ensure compatibility with these new tools, we designed guides covering all NGN PAMs and annotated CBE guides with their NCN sequence context.

To ensure free and easy access to this database, we developed an online portal of the H2M Database v1 (https://human2mouse.com/), which provides user-friendly browsing, visualization, and download of these data. Overall, the H2M Database represents a comprehensive and reliable source for modeling human variants of interest in the mouse genome. We expect to periodically update and expand the H2M database as more human genome sequencing data is collated and analyzed.

### H2M is a multidirectional generator of genetic variant information

In addition to mapping human variants to the mouse genome, H2M is also designed to perform reverse mouse-to-human mapping, as well as other types of functional interspecies modeling on the basis of sequence change effects (**Fig. 4a**). Thus, H2M could integrate experimental data derived from both human cell lines and mouse models with functional interpretation of human variants. We provide three case studies below that broadly illustrate this point and serve as general templates for practical implementation of H2M.

### Case Study 1: Computational modeling of *KIT* human variants with H2M

Multiple studies have shown that the functional similarity of genetic variants between human and mouse is highly dependent on local sequence conservation, even for ortholog pairs with high global conservation^14^. We reasoned that increased flank size similarities suggest higher evolutionary conservation and potential functional significance in the local region. As such, mutations in this region may produce functional effects that are significant and conserved across species.

To illustrate this point, we investigated the human proto-oncogene receptor tyrosine kinase (*KIT*) and its mouse ortholog (*Kit*) as a proof-of-concept. The *KIT* gene encodes a receptor tyrosine kinase that binds the stem cell factor (SCF) ligand and is recurrently mutated or dysregulated in diverse types of human cancer, including gastrointestinal stromal tumors^38^. Both human and mouse orthologs of KIT are composed of extracellular tandem immunoglobulin (Ig) domains, a transmembrane domain, and an intracellular kinase domain (**Fig. 3a-b**)^39^. Clinical studies have identified cancer-associated mutations distributed across all exons of the *KIT* gene; however, recurrent “hotspot” mutations are often located within the transmembrane, juxtamembrane, and kinase domains, suggesting higher functional importance (**Fig. 3c**). Consistent with this, H2M found a significantly higher proportion of human missense mutations within the transmembrane and intracellular kinase domains that can be accurately modeled in the mouse genome (**Fig. 3d**). To investigate whether H2M modeling could also help predict variant pathogenicity, we employed AlphaMissense, which provides human proteome-wide pathogenicity scores for missense mutations based on AlphaFold2 structural predictions^26^, and SIFT (Sorting Intolerant From Tolerant) 4G, which works across multiple species and is based on sequence conservation and amino acid substitution frequencies^40^.

**Figure 3.**
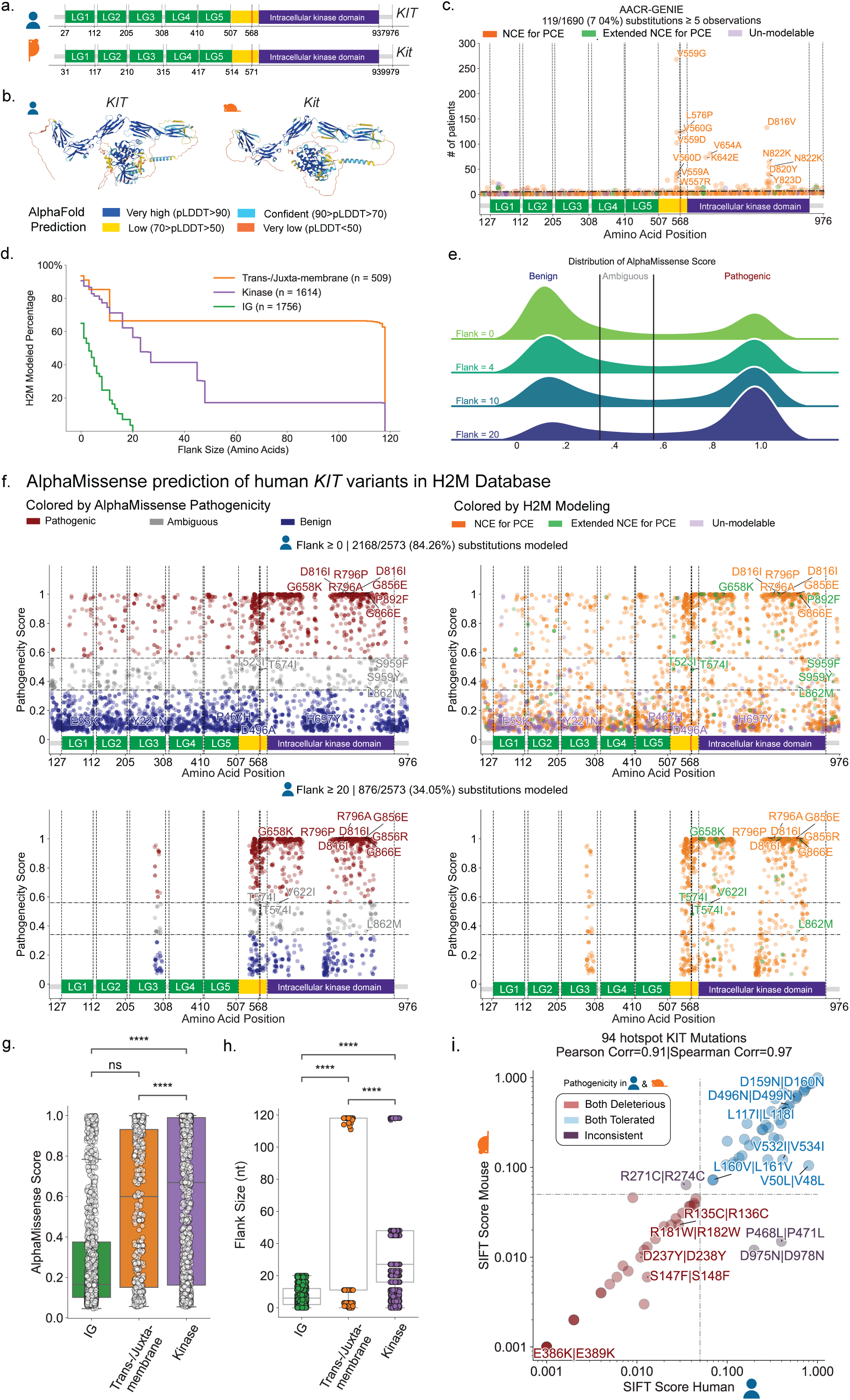
Accurate modeling of human variants with H2M. **a,** Functional domains of the KIT genes (*KIT* in human and *Kit* in mouse), which encode the KIT receptor tyrosine kinase (CD117). LG = Immunoglobulin-like Domain, Yellow = Trans-/Juxta-membrane Domain, Orange = SH2 Binding Domain. Functional domains are labeled according to Uniprot (P10721 · KIT_HUMAN; P05532 · KIT_MOUSE). **b,** AlphaFold predicted structure of human (AF-P10721-F1-v4) and mouse (AF-P05532-F1-v4) KIT protein. **c,** Scatter plot of recurrent frequencies of KIT missense mutations in cancer patients according to AACR-GENIE, colored by H2M modeling. Red dashed line = occurred in 5 patients. H2M-modeling percentage is calculated for unique amino acid substitutions. **d**, Kaplan-Meier Curve visualizing the percentage of human KIT missense mutations that can be modeled by H2M, stratified by functional domain. **e**, Distribution of AlphaMissense scores for all KIT missense mutations in the H2M Database under different thresholds of flank size. Pathogenicity classification: Pathogenic = 0.56-1; Ambiguous = 0.34-0.56; Benign = 0.04-0.34. **f**, Scatter plot of AlphaMissense scores of all KIT missense mutations in H2M Database colored by AlphaMissense pathogenicity (left) and H2M modeling (right), with no flank size limit (top) or flank size ≥ 20 (bottom). H2M-modeling percentage is calculated for unique amino acid substitutions. **g**, Box plot of the AlphaMissense scores of *KIT* variants that can be modeled in mice by H2M, stratified by different functional domains. Statistics shown for t-test of independent samples with Bonferroni correction. **** = p-value ≤ 0.0001, ns = not significant (p-value > 0.05). **h**, Box plot of the flank sizes of *KIT* variants that can be modeled in mice by H2M, stratified by different functional domains. p-values are labeled the same as the (g). **i**, The relationship between SIFT pathogenicity scores for human-mouse mutation pairs in *KIT*, log-scaled. Mutation pairs are selected according to the occurrence of human mutations in AACR-GENIE ≥ 5 patients. Points are labeled by amino acid substitutions in the format of *Human* | *Mouse*. Pathogenicity classification: Deleterious = 0-0.05; Tolerated = 0.05-1. Pearson correlation = 0.91 (p<0.0001); Spearman correlation = 0.97 (p<0.0001).

Increasing the flank size threshold restricted the H2M dictionary to the highly-conserved transmembrane and intracellular kinase domains, which harbor mutations with higher AlphaMissense pathogenicity scores (**Fig. 3e-h, Supplementary Table 6**). In addition, we observed a strong correlation between the SIFT 4G scores of hotspot missense mutation pairs in the *KIT* gene (**Fig. 3i**). These observations imply that increased flank size generally indicates greater evolutionary conservation and functional importance in a region, suggesting that mutations mapping to these types of regions are more likely to have both highly conserved and significant functional impact. Given the clinical importance of mutations in *KIT* and many other therapeutically relevant genes in cancer and other diseases, this case study provides strong support for using H2M to gain insight into gene function, and a roadmap to model clinically-relevant human mutations in the mouse genome.

### Case Study 2: Nominating human neoantigens with H2M

Somatic mutations in cancer cells can generate tumor-specific epitopes that are presented by HLA alleles (H2 in mice), providing ideal targets for cancer immunotherapies^41,42^. However, the widespread deployment of immunotherapies is constrained in part by the limited catalog of targetable neoantigens identified to date, the prohibitive costs of routine whole-genome sequencing in the clinic, and experimental challenges associated with tumor heterogeneity, T-cell cultivation, and immunogenicity assessment^41,43^. Mouse models remain an essential vehicle for *in vivo* discovery and validation of cancer-associated genes and mutations. These models allow for rapid acquisition of primary tumor tissue, deep genome sequencing and mutational calling, and *in vivo* vaccination studies, thereby facilitating the pre-screening and validation of putative neoantigens found in humans^44^. Although a number of studies have identified murine tumor-specific antigens using these types of approaches, the predictive potential and functional conservation of mouse-derived immunogenic mutations in human systems has not been explored. We hypothesized that H2M could be used to systematically predict and functionally map diverse types of immunogenic mutations between different species, including humans and mice.

To test this hypothesis, we first used H2M to determine whether known immunogenic human mutations can produce peptides that are predicted to be recognized and presented by homologous human and mouse MHC molecules, including MHC Class I (MHC-I) and MHC Class II (MHC-II). Of these, MHC-I is known to predominantly present peptides derived from intracellular proteins and lead to induction of cytotoxic T-cell responses^45^. We first retrieved 642 mutation-derived, MHC-I bound neoantigens from the TSNAdb v2.0 online database (**Fig. 4b**)^46^, which has cataloged experimentally-validated tumor-specific neoantigens from the literature. We then used H2M to generate murine versions of the human neoantigens, identifying mouse equivalents for 300 out of 642 neoantigen-producing human mutations. We then used NetMHCpan-4.1 EL, a state-of-art prediction tool trained on mass spectrometry-eluted ligands^45^, to predict MHC-I mutant peptide binding and presentation across the two species. These analyses indicated that > 60% of peptide-pairs are predicted to be presented by at least one MHC allele in both species (**Fig. 4b**). This includes the *EGFR* T790M missense mutation, which is recurrently observed in non-small lung cell cancer patients^47,48^ and known to produce functional T cell epitopes. Importantly, when limiting mouse MHC-I alleles to H2-K^b^ and H2-D^b^, which are expressed by C57BL/6 mice^49^, the same prediction yielded a significant proportion of overlapping immunogenic peptides (**Extended Data** Fig. 3a-b). Together, these analyses validate the utility of H2M to identify functionally conserved neoantigens across species and underscore the value of GEMMs as physiologically-relevant platforms for mechanistic experimental studies of human neoantigens.

**Figure 4.**
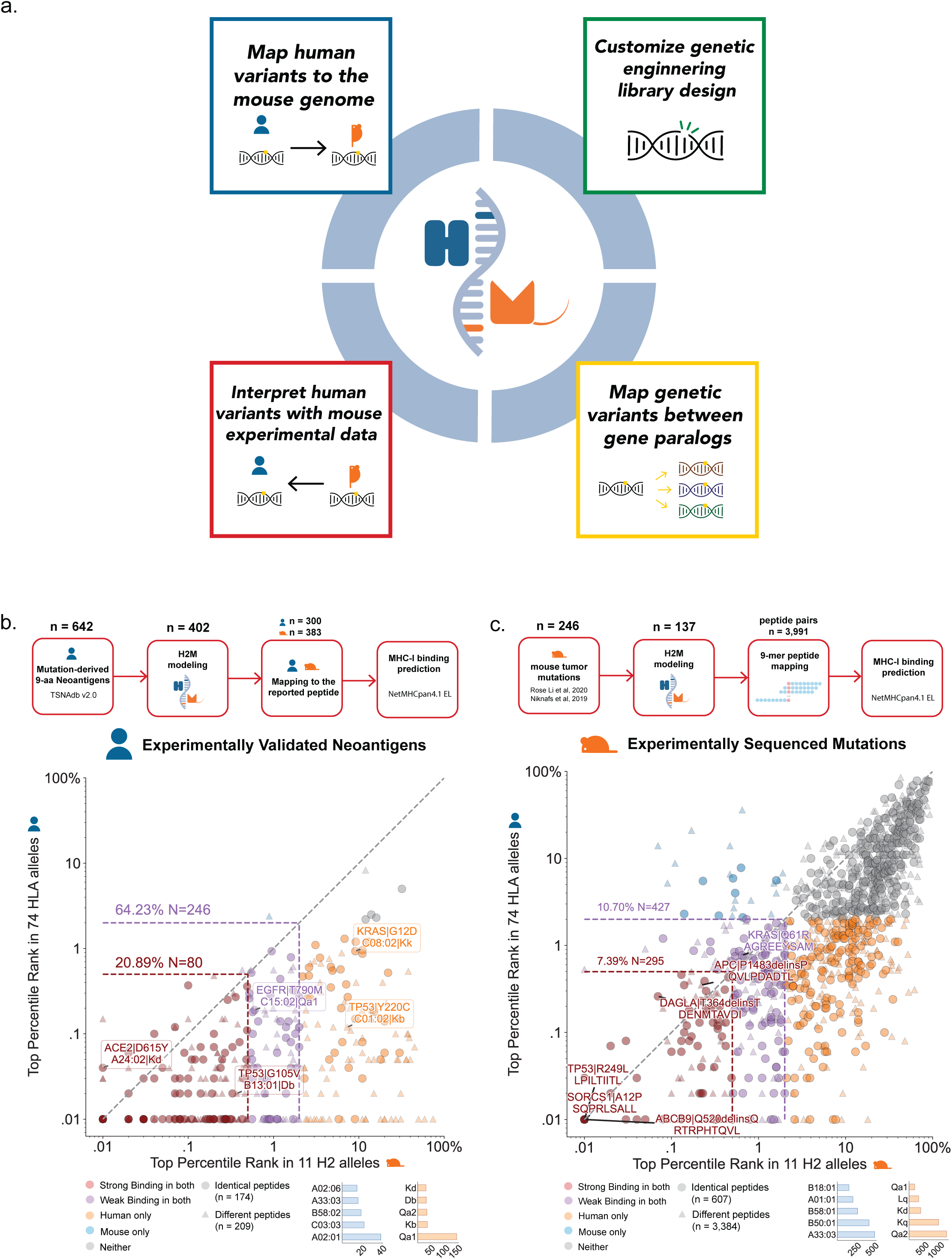
Identification and modeling of conserved immunogenic variants with H2M. **a,** Schematic of potential applications of H2M. In addition to mapping human variants to the mouse genome, H2M is also designed to perform reverse mouse-to-human and paralog-to-paralog mutation mapping, with seamless integration with genome editing library design tools. **b,** Schematic of the generation of human-mouse immunogenic peptide pairs from mutation-derived, experiment-validated human tumor neoantigens, and the relationship of MHC-I binding percentile rank (%Rank) between them. Pearson Correlation = 0.12 (p<0.05), Spearman Correlation = 0.43 (p<0.0001). The top %Rank is selected for each peptide among the predicted set of MHC-I alleles. Top 5 binding alleles are shown in small bar plots for human(blue) and mouse(orange) respectively. Points are colored by binding classifications in both human and mouse. Strong bindings = %Rank < 0.5%, Weak bindings = %Rank < 2%. Circle = Identical 9-mer peptides generated by corresponding mutation in human and mouse; Triangle = Different 9-mer peptides generated by corresponding mutation in human and mouse. **c,** Schematic of the generation of human-mouse peptide pairs from sequenced mutations of mouse tumor models, and the relationship of MHC-I binding %Rank between them. Pearson Correlation = 0.67 (p<0.0001), Spearman Correlation = 0.57 (p<0.0001). Top %Rank selection, binding thresholds, colors, and shapes are the same as in b.

We then tested whether we could simulate the process of discovering potential human MHC-I neoantigens using mouse tumor samples. To do this, we leveraged the species-agnostic nature of H2M to assemble a “M2H” pipeline to analyze a compendium of mutational information obtained by next-generation sequencing of genomic DNA isolated from various types of mouse tumor samples^50,51^ (**Fig. 4c**). This approach identified 246 mutations in mouse protein-coding genes, which are predicted by H2M to generate up to 3,991 neo-peptide pairs in mouse and human cells engineered with equivalent mutations (**Fig. 4c, Extended Data** Fig. 3c**, Supplementary Table 7**). By correlating the rank percentile of MHC allele binding scores, we also identified a significant fraction of murine mutations predicted to be immunogenic in both humans and mice. We also identified a number of synonymous mutations that have been shown to be immunogenic, including the *DAGLA*|T364T-derived FLDENMTAV (IEDB epitope 1889473) and *APC*|P1483P-derived VLPDADTLL (substring of IEDB epitope 472626)^47^ peptides, both of which are recorded as MHC-I T-cell antigens. Many of the non-synonymous candidates we identified have not been recorded previously in the IEDB (Immune Epitope Database), suggesting they may represent tumor-specific neoantigens worth investigating further. High-throughput methods like EpiScan^52^ and TCR-MAP^53^ could be used to interrogate thousands of candidate mutations predicted by H2M to generate immunogenic peptides while sophisticated GEMMs (e.g. *K^b^Strep* mice^54^) could be used for deeper mechanistic studies of high-priority antigens.

Together, these results suggest that functionally-conserved neoantigens could be identified and validated by integrating deep sequencing of mouse tumor-derived genomic DNA, high-throughput genetics/genomics, and functional immunological studies in mice. Some of these neoantigens could be engineered or otherwise installed into the genome of human cells for mechanistic immunotherapeutic studies (e.g. engineered T-cell receptors, vaccines, etc.). Our results establish the potential of integrating cross-species computational analysis with mouse models to discover, predict, and evaluate the immunogenicity of disease-associated mutations to accelerate neoantigen discovery and support personalized medicine efforts.

### Case Study 3: Identifying cognate variants in human gene paralogs with H2M

Paralogous genes play important roles in both normal and disease contexts through functional buffering by independently carrying similarly important roles. Indeed, some paralog pairs are so essential that their combined disruption can trigger potent synthetic sickness or lethality phenotypes^48^. A significant body of work has shown that paralogs play important functional roles in the maintenance of core cellular processes like genome architecture, gene regulation, and mitogenic signaling. Due to their functional redundancy and therapeutic promise, paralogs are currently the subject of intense research.

Identifying functional similarities or interactions between paralogous genes and/or their cognate mutation pairs could advance our fundamental understanding of disease mechanisms and support the development of new targeted therapeutics. Most studies up to date have employed CRISPR-based gene knock-out strategies that lead to complete loss-of-function of one or more paralogs. While powerful, these approaches often ignore the functional consequences of specific types of mutations in each paralog^49^. This is particularly important in cancer and other diseases because paralogs can exhibit a significant degree of functional divergence and specialization, as well as paralog-specific mutational patterns and frequencies depending on the tumor type. Whether different paralogs can exhibit functionally-distinct mutational patterns and how these may impact cancer phenotypes and treatment responses remains unknown.

We reasoned that H2M could enable high-throughput functional interrogation of paralogous mutations by integrating computational searching of mutation equivalents between gene paralogs and across different species with scalable CRISPR-based precision genome editing technologies^35,49,55,56^. To test this idea, we retrieved recurrent cancer-associated single-nucleotide variants from the AACR-GENIE database, filtered them through a literature-curated compendium of human paralog gene pairs^50^, and used H2M to computationally model the mutations in another gene paralog (**Fig. 5a**). This resulted in a catalog composed of 10,221 paralogous mutation pairs (out of 16,225 in total) (**Fig. 5b, Supplementary Table 8**).

**Figure 5.**
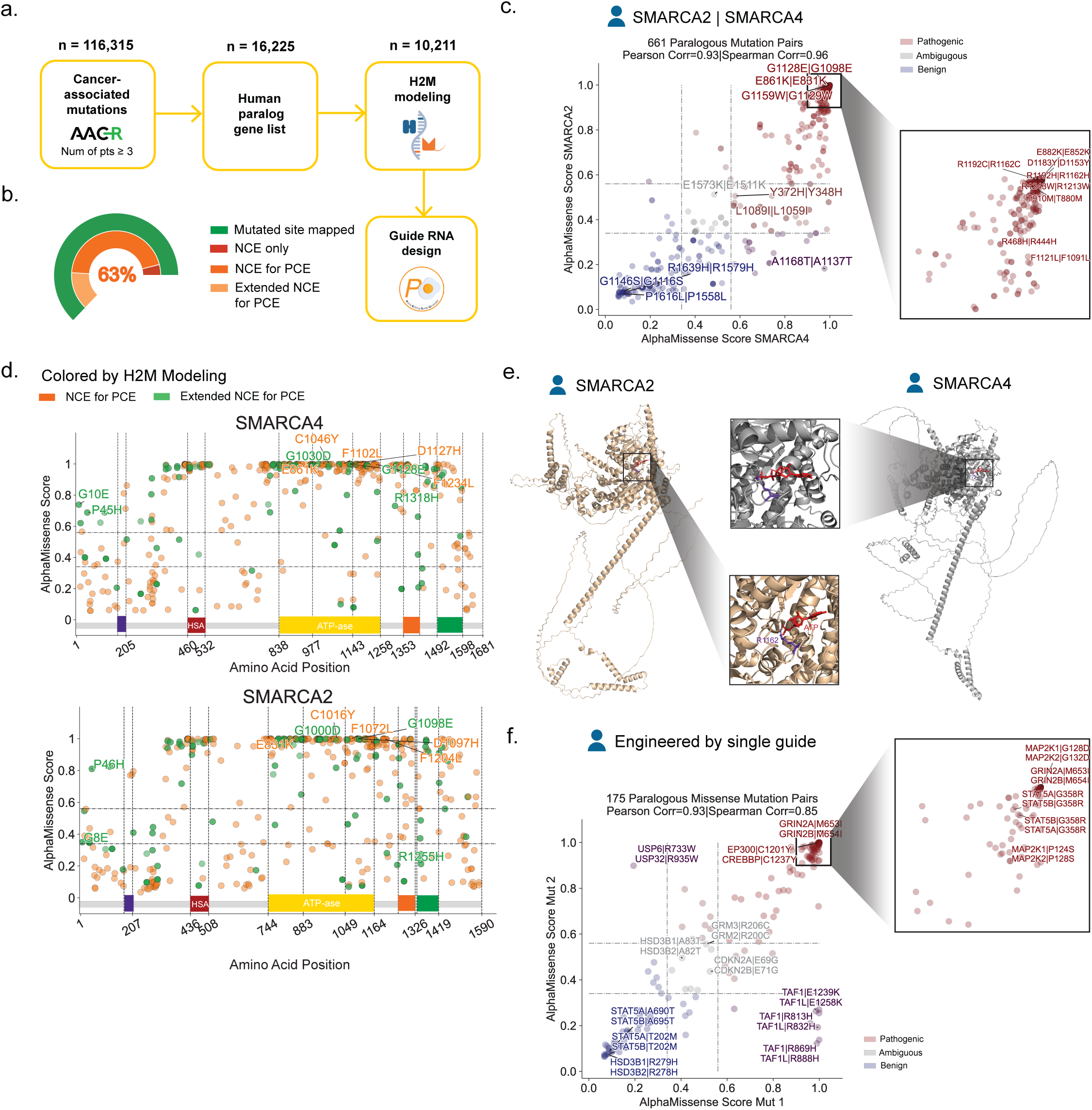
Identification and modeling of conserved variants in paralogs with H2M. **a,** Schematic of modeling human variants across paralogous genes by using H2M. **b,** The percentages of mutations of genes in the H2M Database that can be modeled in the mouse genome, stratified by modeling strategy. **c,** The relationship between AlphaMissense pathogenicity scores for human *SMARCA4* and *SMARCA2* mutation pairs. Points are labeled by amino acid substitutions in the format of *SMARCA4* | *SMARCA2*. Pathogenicity classification: Pathogenic = 0.56-1; Ambiguous = 0.34-0.56; Benign = 0.04-0.34. Pearson correlation = 0.93 (p<0.0001); Spearman correlation = 0.96 (p<0.0001). **d,** Scatter plot of AlphaMissense scores of mapped *SMARCA4* and *SMARCA2* mutations, colored by H2M modeling. No flank size limit. Purple = Gln-Leu-Gln (QLQ) Domain, Red = a helicase SANT-associated (HSA) domain, Yellow = ATP-ase Domain, Orange = SNF2 ATP-coupling (SnAC) domain, Green = Bromo Domain. **e,** AlphaFold 3 predicted ATP (red)-bound structures of human BRG1 (gene: *SMARCA4*) and BRM (gene: *SMARCA2*). H2M-paired residues, BRG1 R1192 and BRM R1162, are highlighted (purple). **f,** The relationship between AlphaMissense pathogenicity scores for human paralogous mutation pairs that can be engineered in parallel with the same base editor and one single guide RNA. Points are labeled by paired genes and the amino acid substitutions. Pathogenicity classification is the same as c. Pearson correlation = 0.85 (p<0.0001); Spearman correlation = 0.84 (p<0.0001).

To illustrate the utility and generalizability of this approach, we focused on the *SMARCA4* and *SMARCA2* paralogs, which respectively encode for the BRG1 and BRM mutually exclusive subunits of the SWI/SNF (BAF) chromatin remodeling complex^57^. Notably, cancer-associated *SMARCA4* mutations are significantly more frequently observed relative to *SMARCA2* mutations, a pattern that also holds true for the *ARID1A-ARID1B* chromatin remodeler paralogs, among others. Supporting functional paralogy, AlphaMissense scores of paired *SMARCA4* and *SMARCA2* variants by using H2M are significantly correlated (**Fig. 5c, Supplementary Table 9**), and the most pathogenic mutations are located in the ATPase and the Helicase/SANT-associated (HSA) domains in both proteins (**Fig. 5d**). Still, the precise functional consequences of each of these variants in *SMARCA4* and *SMARCA2* protein function, or in any of the > 1,000 human genes with known paralogous genomic regions, and how these may vary depending on which paralog is altered is unknown. For instance, the *SMARCA4* R1192C mutation is a statistically significant hotspot and classified as likely oncogenic in OncoKB^58–60^. H2M maps this variant to the homologous *SMARCA2* R1162C mutation and both substitutions are located in the ATP binding pocket (**Fig. 5e**). Both substitutions receive high AlphaMissense pathogenicity scores (**Fig. 5c**), yet clinical observations documenting the effects of R1192C or other *SMARCA2* mutations remain scarce.

To develop a framework to address this problem, we leveraged our mutation catalog to design a base editing library containing > 52,000 unique guide RNAs targeting 4,740 paralogous mutation pairs (**Supplementary Table 8**)^35^. Next, we reasoned that a subset of paralog-targeting gRNAs may be capable of targeting the same paralog pair to engineer the same mutation. In agreement, we found 574 gRNAs targeting 32 genes that can potentially engineer 175 unique paralogous mutation pairs with the same base editor (either cytosine or adenine editor) (**Supplementary Table 9)**. While these gRNAs would otherwise be flagged as off-targeting gRNAs due to targeting more than one homologous region in the genome, H2M is able to integrate cross-species paralogous gene and mutation analyses to identify sequences that can be used for combinatorial paralog mutagenesis. Importantly, these types of mutations are predicted to exhibit a strong functional correlation, as indicated by highly correlated AlphaMissense pathogenicity scores (**Fig. 5f**). Taken together, these results underscore the potential of integrating cross-species genomic analyses with precision genome editing approaches to functionally dissect individual and combinatorial effects of paralogous genes.

## Discussion

Next-generation DNA sequencing technologies have allowed the systematic identification and cataloging of mutations observed in many types of human cancer and other diseases with strong genetic associations. Elucidating the precise functional and mechanistic roles that these mutations play and how these may vary depending on the context remains a highly active area of research. A number of computational and experimental approaches have been developed to tackle this problem; however, little effort has been devoted to improving our ability to perform and interpret systematic cross-species functional studies with the goal of understanding the phenotypic consequences of human genetic variation^41,42^.

To address this problem, we developed H2M, a computational pipeline that systematically models and compares human genetic variation data with other species to both predict their functional consequences and provide scalable precision genome editing tools to test resulting hypotheses. We illustrate the widespread utility of H2M by showing it can perform systematic mouse-to-human and paralog-to-paralog variant mapping coupled to automated prediction and design of bespoke genome engineering libraries that can be deployed for high-throughput genetic studies. Importantly, H2M is not just merely a predictive design tool; instead, we show that the analyses and rich datasets provided by H2M are broadly useful to develop and test new mechanistic hypotheses. For instance, we leveraged H2M to systematically predict and correlate pathogenicity and immunogenicity scores between human-mouse variant pairs, suggesting that variants with similar sequence change effects may also exhibit broad functional conservation between the two species. This presents a testable hypothesis that could be investigated using tailored H2M-derived libraries of gRNAs designed to engineer these conserved human-mouse variant pairs.

The structured framework provided by H2M extends the alignment of genetic information from static sequences to dynamic sequence changes. While the mouse reference genome used by H2M is primarily based on the workhorse C57BL/6 strain, H2M also supports reference genomes from any species, enabling straightforward extension of variant modeling to any other mouse strain or species with available genomes. In doing so, we envision that H2M will open the door for systematic cross-species functional studies of variants (including paralogs) and also inform the development of new physiologically-relevant and genetically-diverse animal models. Such studies would begin to provide critical mechanistic insights into how genetic variation shapes organismal physiology, phenotypic heterogeneity, and disease. The H2M Database (including software package and documentation) can be accessed at https://human2mouse.com.

## Supplementary Tables

Link to supplementary data

**Supplementary Table 1 |** List of homologous genes and canonical transcripts between human and mouse.

**Supplementary Table 2 |** Full H2M Database v1 (AACR-GENIE, COSMIC, and ClinVar) containing >3 million human-mouse mutation pairs.

**Supplementary Table 3 |** Descriptive statistics of H2M Database.

**Supplementary Table 4 |** Database of base editing gRNAs for engineering human and orthologous mouse variants–24,680 gRNAs targeting 2,720 human mutations and 1,892 mouse mutations.

**Supplementary Table 5 |** Database of pegRNAs for engineering human and orthologous mouse variants– 48,255 pegRNAs targeting 4,944 human mutations and 4,707 mouse mutations.

**Supplementary Table 6 |** Pathogenicity prediction of KIT missense mutations in human and mouse.

**Supplementary Table 7 |** MHC-I binding prediction of human-mouse mutational peptide pairs.

**Supplementary Table 8 |** H2H paralogous mutation database.

**Supplementary Table 9 |** SMARCA2/SMARCA4 paralogous mutation pair pathogenicities, and paralogous mutations engineerable by a single base editing gRNA.

## Methods

### H2M Python package

H2M is built as a Python pipeline for Python 3.9-3.12. It is compatible with Mutation Annotation Format (MAF) for both input and output. The online documentation file of H2M is available at https://h2m-public.readthedocs.io.

#### Reference genome and gene annotation

Genome assembly human GRCh37.p13 (GCF_000001405.25) and mouse GRCm39 (GCF_000001635.27) were used as reference genomes for all analyses in this manuscript. Accordingly, GENCODE^18^ comprehensive gene annotations for human GRCh37.p13 ([v19](https://www.gencodegenes.org/human/release_19.html)) and mouse GRCm39 ([vM33](https://ftp.ebi.ac.uk/pub/databases/gencode/Gencode_mouse/release_M33/)) genomes were used for coordinating transcripts in each respective genome.

#### Retrieving ortholog pairs, transcripts, and canonical transcripts

A list of human-mouse ortholog gene pairs was generated by integrating orthologous annotations from the Ensembl and MGI (Mouse Genome Informatics) databases. A list of human genes (protein coding genes, mitochondrial genes, genetic regulatory elements, etc.) and their murine orthologs were retrieved using the Ensembl BioMart data mining tool (implemented with pybiomart) and combined with all entries downloaded from MGI^21–24^ [Vertebrate Homology Table](https://www.informatics.jax.org/downloads/reports/index.html#homology). The complete list of transcripts, including unique canonical transcripts for each human or mouse gene, were retrieved via Ensembl API.

#### Sequence alignment

If a mutation overlaps with the stop codon, the human-mouse pairwise sequence alignment is based on the location of the stop codon; otherwise, the Needleman–Wunsch algorithm is used via the Bio.pairwise2 module from biopython 1.81^51^ without gap penalty. Identical characters have a score of 1 (otherwise 0). The alignment with the highest score is selected. If multiple alignments exist with the highest score, the first one returned is selected. Peptides are aligned for the modeling of coding mutations while transcripts are aligned for non-coding mutations.

### Generation of H2M Database

Human clinically-observed variants were curated from [AACR Project GENIE](https://www.aacr.org/professionals/research/aacr-project-genie/) (syn7222066, v15.0, released Apr 8, 2024)^23^, [COSMIC Census Genes Mutations](https://cancer.sanger.ac.uk/cosmic) (v99, released Nov 28, 2023)^33^, and [ClinVar](https://www.ncbi.nlm.nih.gov/clinvar/) (retrieved Feb 6, 2024)^34^. GENIE data was offered only in the GRCh37 version and MAF format. For COSMIC and ClinVar data, the GRCh37 versions were selected, and the VCF files were converted to the MAF format by using H2M. For GENIE and COSMIC data, originally-provided human transcripts were used. For ClinVar data, Ensembl Canonical human transcripts were used. Some of the gene symbols were manually checked and renamed to match the list in H2M due to the usage of gene name aliases.

H2M 1.0.3 was then used to generate modeling of murine equivalents. Homologous gene(s) of a given human gene were all included. Ensembl Canonical murine transcripts were used. Up to 5 extended NCE for PCE alternative modeling were included for each human mutation. The human-mouse mutation dictionary database is provided in MAF format and available for online browsing at https://human2mouse.com.

### *In-silico* library design for precision genome editing

A subset of mutations from the H2M database that originated from the AACR database and had an observation count > 20 was selected for library construction. Guide RNAs were designed by using PEGG^35^. For both base editing and prime editing libraries, NG PAM sequences were considered. For base editing, both cytosine base editors (CBEs) and adenine base editors (ABEs) were considered. For prime editing, up to 5 guide RNAs were designed for each mutation, which were ranked and selected by the “RF_Score,” a random forest predictor of pegRNA activity.

### Genome coordinate conversion

The [UCSC Lift Genome Annotations](https://genome.ucsc.edu/cgi-bin/hgLiftOver) website tool was used to convert H2M-derived GRCm39 mutations to GRCm38.

### Evaluation of missense mutation pathogenicity with AlphaMissense and SIFT

AlphaMissense scores for all human amino acid substitutions, which were labeled by the Uniprot IDs of the proteins, were downloaded from AlphaMissense^26^ Google Cloud page. Based on the AlphaMissense score, the classification for pathogenicity was: Pathogenic (0.56-1); Ambiguous (0.34-0.56); Benign (0.04-0.34).

SIFT 4G is a faster version of SIFT, which also provides SIFT predictions for more organisms^40^. The SIFT 4G database for human (GRCh37) and mouse (GRCm38), as well as the SIFT 4G annotator software, were downloaded from the [SIFT website](https://sift.bii.a-star.edu.sg/sift4g/AnnotateVariants.html). 94 *KIT* missense mutations generated by single-nucleotide alterations were classified as “hotspot” mutations based on their occurrence in more than 5 cancer patients in AACR-GENIE. Selected *KIT* hotspot missense mutations and their mouse equivalents were exported to VCF files in order to be annotated with SIFT 4G scores by using SIFT 4G annotator software. Based on the SIFT 4G score, the classification for pathogenicity is: Deleterious (0-0.05); Tolerated (0.05-1).

### Generation of human-mouse peptide pairs for immunogenic prediction

#### From human validated peptides

Human immunogenic missense mutations and specific MHC-I binding peptides (9 amino acids long) generated from their corresponding mutant proteins, were curated using [TSNAdb v2.0](https://pgx.zju.edu.cn/tsnadb/) ^46^. H2M was then used to map immunogenic mutations based on the same peptide change effect. Following this, corresponding murine neo-peptides were mapped to the reported human ones, based on the identical relative position of the mutated amino acid in the neo-peptide.

#### From mouse tumor mutations

Mouse mutational data was curated from supplemental materials obtained from previously published studies (Rose Li et al, 2020 ^58^ and Niknafs et al, 2019 ^59^). Up to 9 of all the possible 9-mer neo-peptides derived from each mutation were generated. Following this, H2M was used to map murine mutations based on the corresponding peptide change effect in the human genome. Corresponding human neo-peptides were mapped to pre-generated human ones based on identical relative position of the mutated amino acid in the neo-peptide.

### MHC-I binding prediction of mutation-derived neoantigens using NetMHCpan4.1 EL

Prediction tasks were performed online by using NetMHCpan4.1 EL^45^ (recommended epitope predictor-2023.09), available at http://tools.immuneepitope.org/mhci/. Peptides were input in FASTA format. The length of binding peptides was set to 9 amino acids for both human and mouse. A set of MHC-I alleles was selected and the top percentile rank of each peptide was selected in the analysis.

NetMHCpan methods inform if a sequence is a strong MHC binder (SB) or a weak MHC binder (WB) based on the percentile and score. Percentile rank (%Rank) of a query sequence was computed by comparing its prediction score to a distribution of prediction scores for the MHC in question, estimated from a set of random natural peptides. For MHC-I, %Rank < 0.5% and %Rank < 2% thresholds were considered for detecting SBs or WBs.

Selected Mouse MHC alleles list (11 in total): (H-2-) Db, Dd, Dq, Kb, Kd, Kk, Kq, Ld, Lq, Qa1, Qa2.

Selected Human MHC alleles list with frequent occurrence (74 in total): (HLA*) A01:01, A02:01, A02:06, A03:01, A11:01, A23:01, A24:02, A25:01, A26:01, A29:02, A30:01, A30:02, A31:01, A32:01, A33:03, B07:02, B08:01, B13:01, B13:02, B14:02, B15:01, B15:02, B15:25, B18:01, B27:02, B27:05, B35:01, B35:03, B37:01, B38:01, B39:01, B40:01, B40:02, B44:02, B44:03, B46:01, B48:01, B49:01, B50:01, B51:01, B52:01, B53:01, B55:01, B56:01, B57:01, B58:01, B58:02, C01:02, C02:02, C02:09, C03:02, C03:03, C03:04, C04:01, C05:01, C06:02, C07:01, C07:02, C07:04, C08:01, C08:02, C12:02, C12:03, C14:02, C15:02, C16:01, C17:01, E01:01, E01:03, G01:01, G01:02, G01:03, G01:04, G01:06.

### Data availability

Part of the public data used in this study, including reference genome, gene annotations, and public datasets of human variants, as well as figure-related data, including H2M Database, paralogous mutation pairs, as well as the corresponding genome editing libraries, is available in the following <u>Dropbox</u> folder. The H2M Database, including the human-mouse mutation dictionary, and the genome editing guide RNA library, is available for online browsing at http://human2mouse.com and for download in the associated Dropbox folder.

### Code availability

The newest version of H2M (1.0.3) is freely available on PyPI at https://pypi.org/project/bioh2m/ and on GitHub at https://github.com/kexindon/h2m-public. A Tutorial for using H2M is also provided in the GitHub repository, in addition to all analysis scripts and codes for the generation of figures. Further documentation and installation instructions for PEGG are available at https://h2m-public.readthedocs.io.

## Acknowledgements

We thank all members of the Sánchez-Rivera laboratory for feedback and support. We thank B. Ding and F. Yue for assistance in developing the H2M web portal. We thank Y. Soto-Feliciano, N. Mathey-Andrews, M. T. Hemann, and J. S. Weissman for scientific discussions and overall support. We thank Claire Glickman for laboratory management support and Jamie Rothman for administrative support. We also thank the Koch Institute’s Robert A. Swanson (1969) Biotechnology Center for technical support, especially the Barbara K. Ostrom (1978) Bioinformatics Facility. Work in the Sánchez-Rivera laboratory is supported by the Howard Hughes Medical Institute (Hanna Gray Fellowship, GT15656), the V Foundation for Cancer Research (V2022-028), NCI Cancer Center Support Grant P30-CA014051, the Virginia and D.K. Ludwig Fund for Cancer Research, Koch Institute Frontier Research Program, the Casey and Family Foundation Cancer Research Fund, the Michael (1957) and Inara Erdei Fund, the MIT Research Support Committee, the Upstage Lung Cancer Foundation, and a Traditional Project Award from the Bridge Project, a partnership between the Koch Institute for Integrative Cancer Research at MIT and the Dana-Farber/Harvard Cancer Center. S.I.G. is supported by NIH T32GM136540 from the NIH/NIGMS. S.I.G. is also supported by a MIT School of Science Fellowship in Cancer Research.

## Author contributions

K.D., S.I.G., and F.J.S.R. conceived the project and wrote the manuscript. K.D. developed H2M, performed all computational analyses, assembled all figures, and conceptualized the H2M web portal. S.I.G. provided conceptual and technical support, and additional day-to-day mentorship to K.D. M.L. optimized the H2M 2.0 web portal and the underlying associated SQL database, as well as optimized its backend performance. F.J.S.R. supervised the work and secured funding.

## Ethics declarations

F.J.S.R. has consulted for Repare Therapeutics and Ono Pharma. The remaining authors declare no competing interests.

---

**Extended Data Figure 1.**
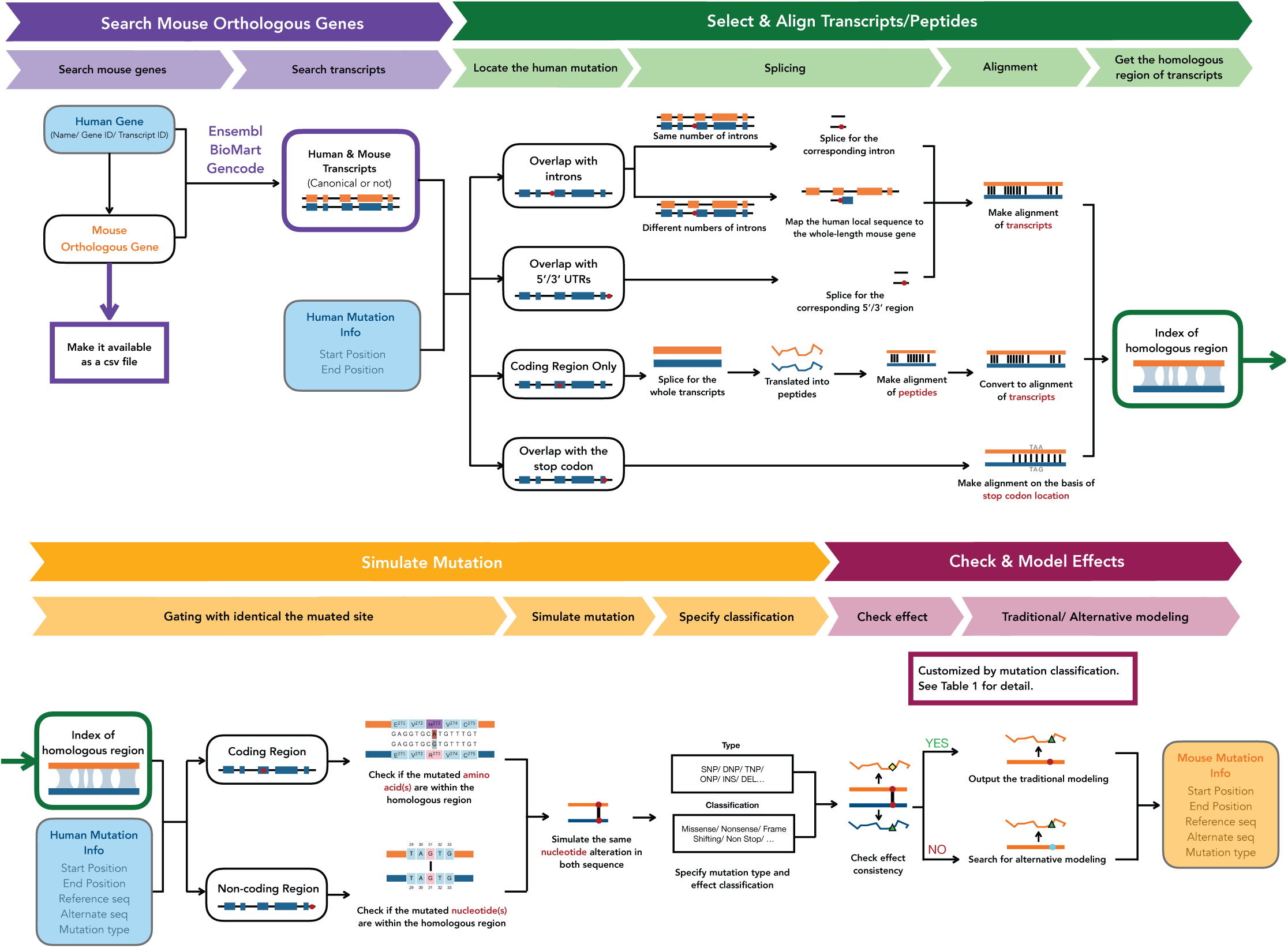
Detailed schematic of H2M.

**Extended Data Figure 2.**
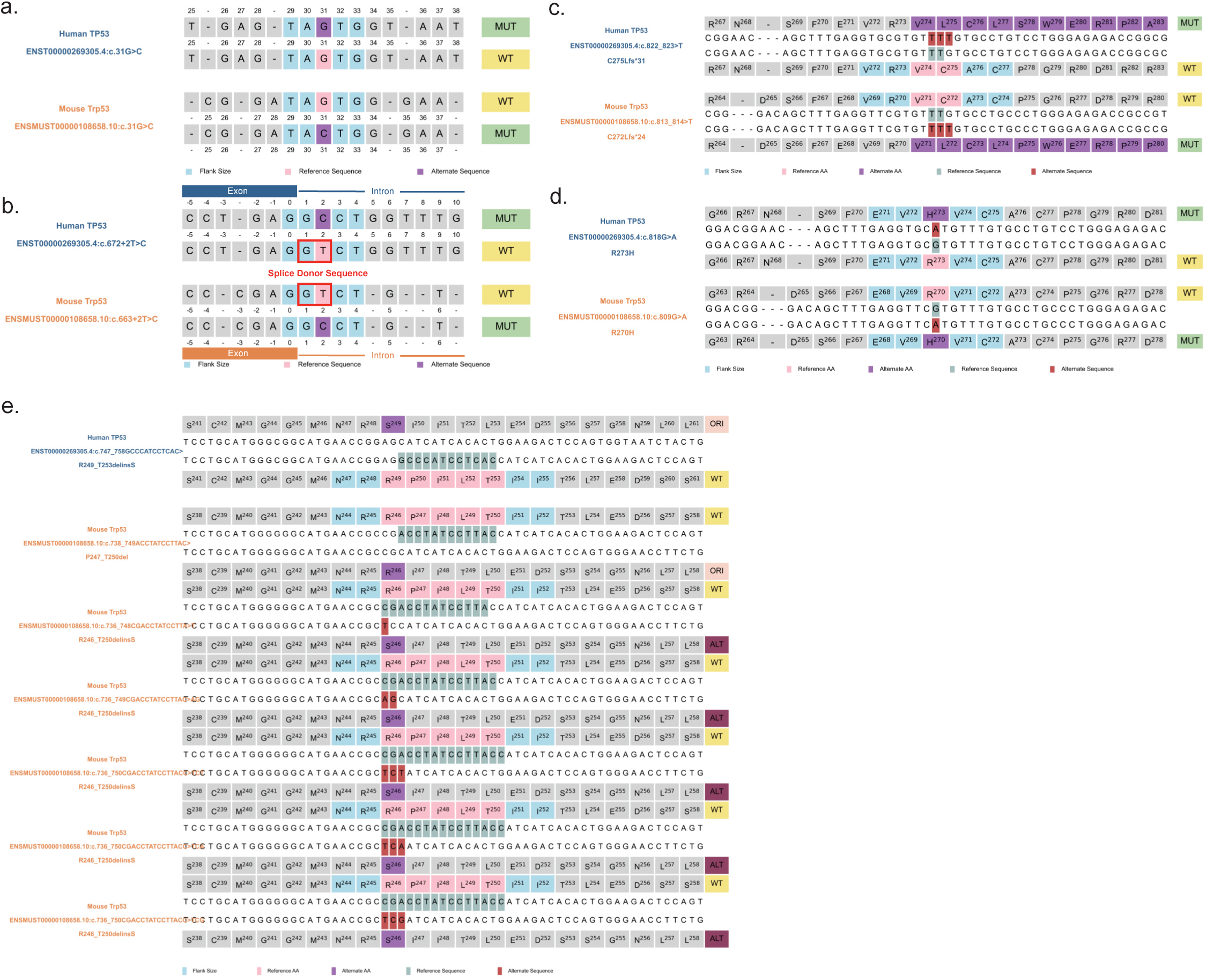
Visualization examples of p53 mutations generated by H2M. **a,** NCE-only modeling of a single-nucleotide variation in a sample intron. **b,** NCE-only modeling of an insertion in a sample splice site. **c,** NCE-only modeling of a frame-shifting insertion in a sample coding region. **d,** NCE for PCE modeling of a missense mutation in a sample coding region. **e,** Extended NCE for PCE modeling of an in-frame deletion in a sample coding region.

**Extended Data Figure 3.**
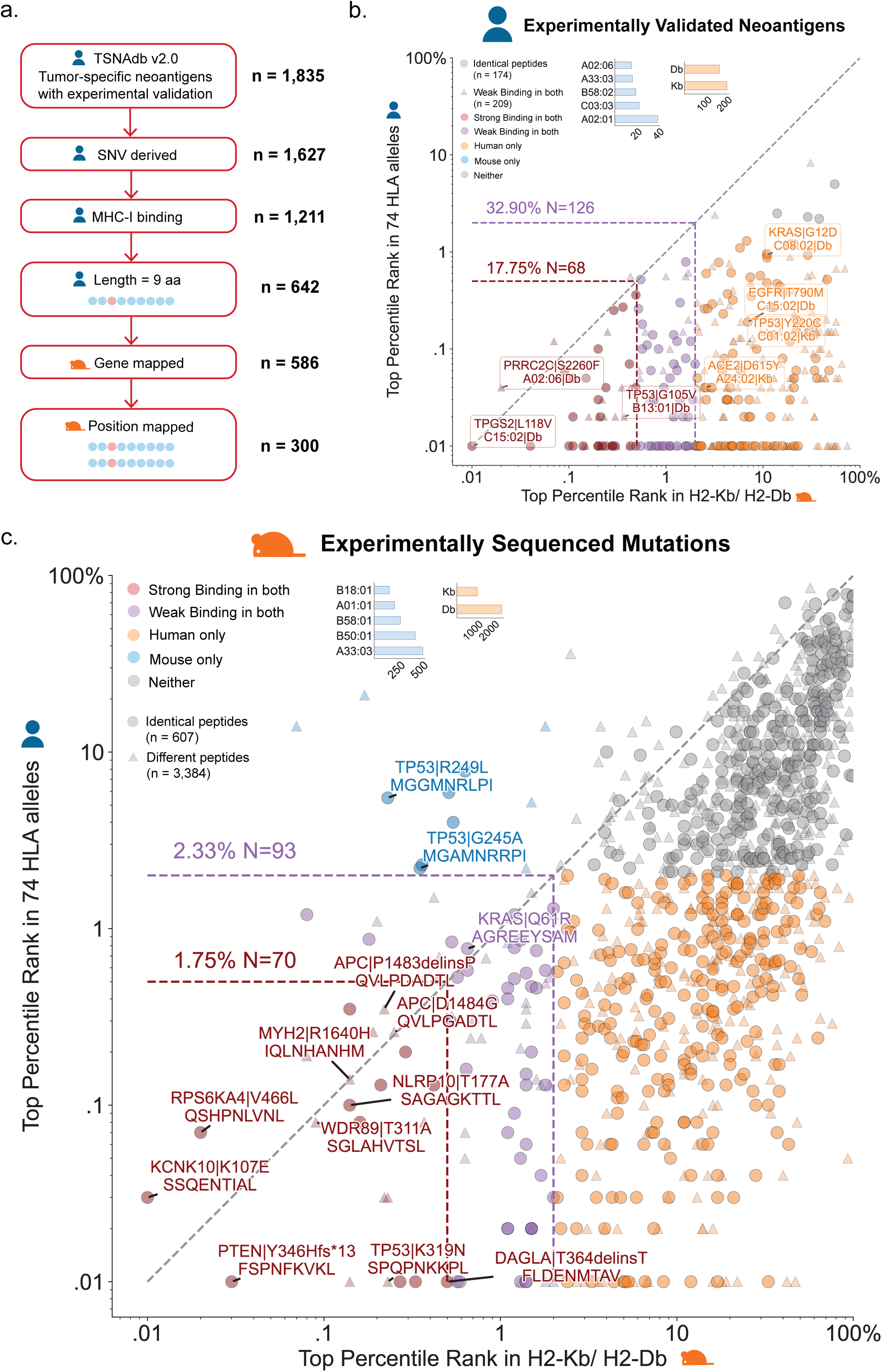
Functional conservation of immunogenic human-mouse peptide pairs. **a,** Detailed schematic of the generation of human-mouse immunogenic peptide pairs from validated human ones, and **b,** the relationship of MHC-I binding %Rank between them. Pearson Correlation = 0.17 (p<0.05), Spearman Correlation = 0.25 (p<0.0001). For human peptides, the top %Rank is selected for each peptide among the set of 74 MHC-I alleles (top 5 counts shown in blue bars). For mouse peptides, the top %Rank is selected for each peptide among C57BL/6-strain expressed MHC-I alleles, H2-Kb and H2-Db (counts shown in orange bars). Strong bindings = %Rank < 0.5%, Weak bindings = %Rank < 2%. **c,** The relationship of MHC-I binding %Rank between human-mouse peptide pairs derived from mouse cancer-associated somatic mutations. Pearson Correlation = 0.49 (p<0.0001), Spearman Correlation = 0.54 (p<0.0001). Top %Rank selection, binding thresholds, colors, and shapes are the same as b.

